# Detection and localization of conspecifics in ghost knifefish are influenced by the relationship between the spatial organization of receptors and signals

**DOI:** 10.1101/2023.07.20.549925

**Authors:** Keshav L Ramachandra, Oak E Milam, Federico Pedraja, Jenna Cornett, Gary Marsat

**Affiliations:** Department of Biology, West Virginia University, Morgantown, USA; Department of Neuroscience, West Virginia University, Morgantown, USA; Department of Neuroscience, Zuckerman Mind Brain Behavior Institute, Columbia University, New York, USA

## Abstract

The detection and localization of signals rely on arrays of receptors, and their spatial organization plays a key role in determining the accuracy of the system. Weakly electric ghost knifefish rely on a distributed array of electroreceptors to detect spatially diffuse electric signals from conspecifics. While we know that spatial resolution for small objects, such as prey, is enhanced near the head due to a high receptor density, it is not clear how receptor organization influences the processing of global and diffuse signals from conspecifics. Using spatially realistic modeling, we quantified how receptor density influences detection and localization accuracy for conspecific signals across varying distances.

Our main result demonstrates that receptor density markedly enhances detection accuracy in frontal regions at intermediate distances (35-50 cm) yet surprisingly contributes minimally to improving localization accuracy. This highlights a fundamental principle: receptor convergence primarily benefits signal detection when dealing with spatially diffuse stimuli, even though higher receptor density can enhance localization accuracy for spatially delineated signals. Our findings extend beyond the electrosensory modality, drawing parallels with other sensory systems, and offer broader insights into spatial processing principles.

## INTRODUCTION

Accurate detection and localization of sensory signals is fundamental to animal behavior, influencing predator avoidance, mating decisions, and resource acquisition. Signal detection and localization typically rely on an array of sensory receptors; their sensitivity, number, and spatial organization play a key role in setting the accuracy of the system. Therefore, understanding how receptor arrays are spatially structured to optimize these sensory tasks remains a central question in sensory neuroscience. For example, visual resolution is enhanced by the high receptor density in the retina’s fovea, sound localization is largely enabled by the binaural configuration of the auditory system, and moths can detect the presence of just a few pheromone molecules due to the number and convergence of olfactory receptors (Ashida and Carr, 2011; Provis et al., 2013; Rospars et al., 2014). While the spatial structure and size of the receptor array influence the system’s sensitivity, it is not always clear how the configuration of the receptors relates to detection accuracy versus localization. This is particularly true for diffuse signals that affect most of the sensory array, such as a luminosity gradient in the environment, a distant heat source, or the Earth’s magnetic field.

The electrosensory system in fish is a remarkable example of sensitivity. In ghost knifefish, in particular, survival depends on navigating, detecting prey, and communicating through this active sense. They generate a constant weak electric field with their electric organ (EO), and any distortions of this field by preys or objects in their environment are picked up by an array of receptors covering the skin of the fish (Nelson et al., 1997; Pedraja et al., 2014). Distortions from an object or prey will impact only a spatially defined subset of receptors on the corresponding portion of skin onto which the electric image (EI) of the object is projected. By activating the corresponding portions of the topographic maps higher in the sensory system, spatial information is encoded in a sort of labeled-line code reminiscent of the way the visual or somatosensory system is organized. Similar to these modalities (i.e., fovea of the retina), regions of higher receptor density towards the head and snout of the fish provide a higher spatial resolution particularly useful in the last stage of prey capture, as the target approaches the fish’s mouth (MacIver et al., 2001; Nelson and Maciver, 1999). They also detect and communicate with one another through this electric sense (Allen & Marsat, 2018; Knudsen, 1975; Petzold et al., 2016). The ongoing electric organ discharge (EOD) is a spatially diffuse signal that can reach globally all the receptors of the other fish’s body. Localization of such signals would thus have to rely on differences in the signal strength at different input locations, similar to the way the auditory system compares binaural inputs to localize sound sources (see Milam et al., 2019, for review). While it is clear that the high rostral density of receptors can enhance the spatial accuracy for objects, it is not clear how it influences the processing of diffuse and global signals from another fish’s EOD. More specifically, we are interested in determining how the spatial organization and density of receptors interact with the spatial structure of EOD signals to influence the detection and localization of conspecific signals.

The EOD generated by the long EO located in the caudal 2/3 of the fish can be approximated as a dipole whose polarity switches during each EOD cycle (Rasnow et al., 1993). The resulting signal is a quasi-sinusoidal output with frequencies between 500 Hz and 1000 Hz (in *A. leptorhynchus*, the focal species in this paper; (Zupanc and Maler, 1993). Although weak, this signal will travel several tens of cm and will permit the long-range detection of the conspecific (Pedraja et al., 2016). Behavior studies demonstrated that frequent interactions occur at distances of 30 cm or less, but there is evidence from field studies that these fish might be able to detect and navigate toward one another at distances in the 1-meter range (this upper limit has not been quantified systematically; (Henninger et al., 2018; Stamper et al., 2012; Stamper et al., 2013; Zupanc et al., 2006). These distant signals will reach the receptors with low intensity as the strength of electric signals decreases exponentially with distance. The signal from a distant fish will combine with the signal of the fish’s own EOD, resulting in a combined electric field with sinusoidal amplitude modulations designated as the beat. If the distant fish’s signal reaches the focal fish with an amplitude 1/10^th^ the strength of the self-generated EOD, this beat modulation will have an amplitude of 1/10^th^ the undisturbed EOD. These weak beat contrasts are the signals that must be encoded to detect a conspecific, and differences in contrast at receptors’ locations across the fish’s body constitute the localization cues. The mathematical framework to estimate the strength and structure of the electric field during social interactions has been detailed in previous studies (Caputi and Budelli, 2006; Castello et al., 2000; Gómez-Sena et al., 2014; Kelly et al., 2008). It can be used to quantify the strength of the signal as it reaches each receptor to obtain a complete characterization of the sensory input structure due to an approaching conspecific.

Ghost knifefish possess several types of electroreceptors; we focus here on p-unit tuberous receptors, which constitute the vast majority of electroreceptors, are tuned specifically to encode conspecific signals, and are responsible for encoding the amplitude of these beat contrasts (Bennett et al., 1989). Previous estimates of p-units receptor density range from 9-15 per mm^2^ on the head region to 0.6-3.4 over the trunk area (Carr et al., 1982). A thorough quantification of receptor density as a function of dorso-ventral/rostro-caudal location in *A. leptorhynchus* is not yet available. Receptor sensitivity and response properties have been extensively studied, and several neural models of p-units are available (Benda et al., 2005; Chacron et al., 2005; Goense and Ratnam, 2003; Gussin et al., 2007; Nelson et al., 1997; Ratnam and Nelson, 2000). This large population of several thousand receptors converges to the primary sensory area in the hindbrain, the electrosensory lateral line lobe, and the information is then transmitted down the sensory pathway (Lannoo et al., 1989; Maler et al., 1991). To estimate the information carried by the population of receptors about realistic signals, two key elements must be considered. First, we must consider the response properties of the receptors, including their sensitivity and noise levels, as well as their heterogeneity across the population. Second, we must consider the structure of the stimuli and how signals from different locations will result in input strengths that vary across receptor positions.

In this paper, we aim to clarify how spatially realistic signals from conspecifics are encoded by the population of electroreceptors. Our approach includes using a model of the fish’s electric field to quantify the EI strength at each receptor location. This input drives a model of the p-unit population consisting of heterogeneous leaky-integrate-and-fire units calibrated based on the extensively documented properties of this population. We then use a decoding analysis to estimate the information that can be extracted from the receptor population and provide a conservative estimate of the expected detection and localization accuracy. We specifically hypothesize that the high density of receptors rostrally will enhance detection and localization accuracy, particularly in the frontal quadrant. We tested this hypothesis by altering the structure and density of the receptor population and confirming that detection accuracy depends on receptor density. Surprisingly, we demonstrate that localization is relatively less influenced by receptor density, and that high rostral density does not enhance localization accuracy for frontal azimuth. Our results reveal the intricate relationship between the spatial structure of signals and the spatial organization of sensory receptors, highlighting broader sensory processing strategies relevant across modalities and taxa.

## METHODS

### Quantification of electroreceptor distribution

Weakly electric brown ghost knifefish, *Apteronotus leptorhynchus*, were obtained from a tropical fish supplier (Segrest Farms, FL, USA). Fish care and use were approved by West Virginia University IACUC.

Fish were euthanized, then fixed in a 50mL aliquot containing a 40mL solution of 4% paraformaldehyde and preserved for up to 7 days. After the tissues were completely fixed, 5mg of eosin Y was added to the aliquot containing the preserved fish. Stained fish were analyzed under a fluorescent microscope with a light wavelength of 530nm. A stereotaxic system was used to move the fish and place a 1 mm² sampling grid at different locations along the fish’s body. Tuberous receptors inside the grid were visually identified and counted. We collected 581 samples from 18 fish measuring, on average, 14 cm in length.

A coarse 3D mesh model of the fish was created using Maya 2019 (Autodesk, Inc.) based on average measurements of our fish and images of the rostral, dorsal, and lateral profiles. The quadrangle mesh model has 218 planar faces. For each face, the corresponding measured receptor densities were averaged. Average receptor densities for each face were mapped along a rostro-caudal and dorsal-ventral plane. A 3-dimensional 5^th^-degree polynomial was fitted to obtain a smooth, interpolated estimate of receptor density as a function of body location. Using a mesh model with triangular faces (the same used for EI calculation; see below), we randomly generated receptor locations for each face according to our density function.

### EI model

The electric image model used in this study was based on the established methods developed by Caputi and Budelli (Caputi and Budelli, 1995; Caputi and Budelli, 2006; Caputi et al., 1998) and implemented using software developed by Rother (Rother et al., 2003). More details on the model can be found in the following publications, and a brief description is provided here. The EI model requires the creation of a reconstruction of the geometry and electrical properties of the fish bodies and their placement in the surrounding water. This information is used to calculate the transcutaneous voltage at specific nodes along the skin of the fish. The model makes the following assumptions:

1. All the media are ohmic. Therefore,

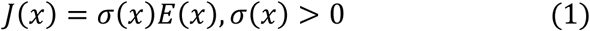

Where J(x) is the current density at point x and E(x) is the electric field at the same point.
2. There are no capacitive effects, so at no point in space is there an accumulation of charge.

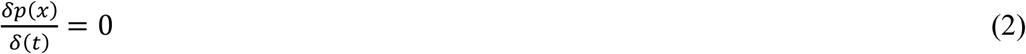
3. The model is an electrostatic approximation (Bacher, 1983)
4. The fish and other objects that make up the environment are immersed in an infinite water medium. Each object in the environment is covered by a thin resistive layer (the skin in the case of the fish), which can be homogeneous or heterogeneous.

The model is based on the charge density equation, which, under the above assumptions, implies that the charge generated by the sources f(x) is equal to the charge diffusion:

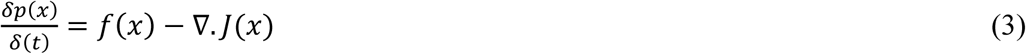

Combining equations 1 and 2, we get

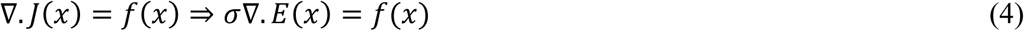

The electric field E(x) can be expressed as *E*(*x*) = −∇*φ* therefore:

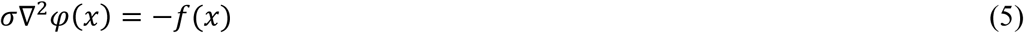

Equation (5) is a partial differential equation known as the Poisson equation, which can be solved for every point in space, in this case, the fish boundaries, using the boundary element method (BEM) as proposed by Assad (Assad and Bower, 1997). The method determines the boundary conditions by solving a linear system of M•N equations for M poles and N nodes, with the unknown variables being the transepithelial current density and voltage at each node (Pedraja et al., 2014). The shape of the 3D fish mesh model consists of 49 ellipses composed of 17 nodes each (i.e., 835 nodes), defining 1,666 triangular faces between the nodes (Rother et al., 2003).

The size of the fish was kept constant at 14 cm in length in this paper, the water conductivity was set to 300 µS, and the skin and internal conductivity of the fish were 100 and 10,000 µS, respectively. The two poles for each fish were positioned 9.3 and 10.5 cm from the rostral tip of our 14 cm fish. We use the middle of this dipole (i.e., the center of our “electric organ”) to define the position from which the signal originates.

### Calculations of signal strength at receptor locations

We calculate the transdermal voltage when only the focal fish is present (*Vf*) or when both focal and sender fish are present (*Vfs*), in which case the peak voltage is determined at the top of the beat cycle (EODs in phase). The strength of the EI caused by the sender fish at each node *i* on the surface of the focal fish was defined as a contrast *c*:

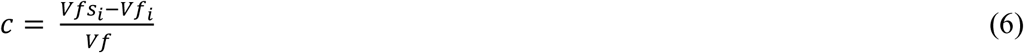

The contrast value for receptor location within each triangular face was interpolated from the values at the nodes that define the face using a barycentric coordinates system.

Considering a triangle with values *N* at the nodes (vertices) and coordinates *X, Y, Z*. A receptor inside the triangle will have a contrast value *R* according to the weighted value of the nodes. The weights *W* can be found by solving:

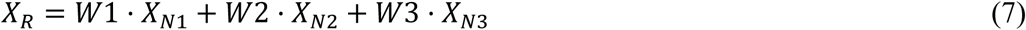

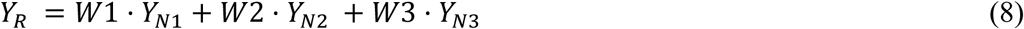

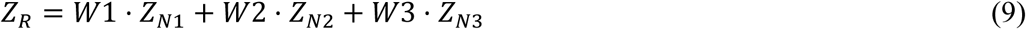

Contrast at the receptor location is calculated as:

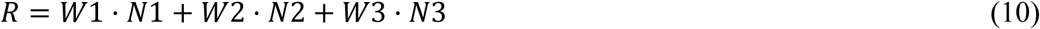

Although the EI model was thoroughly calibrated based on experimental recordings on actual fish (Pedraja et al., 2014; Pedraja et al., 2016), we performed transcutaneous recordings on three pairs of fish and verified that the transdermal voltage contrast values that we calculated correspond to the range of values that can be measured experimentally (data not shown).

### Receptor Modelling

We used a leaky integrate-and-fire (LIF) framework to model the receptors. The model includes noise σ and adaptation current with conductance *g_α_*, reversal potential *E_α_*, and is driven by an input *I*. The membrane voltage is calculated as:

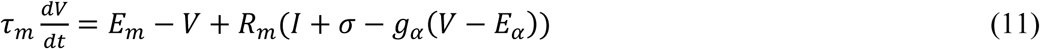

The initial parameters of the model were based on existing models (Benda et al., 2005; Carlson and Kawasaki, 2006; Chacron et al., 2001; Nelson et al., 1997). Particularly, the noise was the product of a strength variable *A_σ_* and a random process (specifically, two Ornstein– Uhlenbeck processes (refer to Chacron et al., 2001, for details). Adaptation current *α* was adjusted to match the time course of adaptation described experimentally (Benda et al., 2005). Conductance *g_α_* is augmented by *Δ_α_* after each spike and decays with time constant *τ_α_*. When the membrane voltage reaches the threshold *V_T_* it is reset to *V_R_* and kept constant during a refractory period *t_R_*. The input *I* to these p-unit model neurons replicates the input they would receive from receptor cells during social interactions. A sinusoidal EOD carrier signal with amplitude *A_EOD_* was created with a frequency of 1000 Hz (the upper range of the naturally occurring EOD frequencies in this species was used for convenience) and modulated with a sinusoidal amplitude modulation (the “beat”) of 30 Hz (other AM envelopes are used to validate our model, see Supplementary Material). This AM envelope signal was adjusted to a specific contrast as specified in the Results. A contrast of 0% indicates that the baseline EOD is unmodulated, whereas a beat contrast of 100% causes the EOD amplitude to be zero during the trough and twice the baseline EOD amplitude at the peak of the beat. To emulate the current direction and sensitivity as the signal passes through the receptor cells before it reaches the p-unit neurons, the modulated EOD is halfwave rectified after a baseline bias *β* was subtracted.

Parameters were adjusted to create a prototypical neuron with response properties matched to published data (Bastian, 1981; Benda et al., 2005; Chacron et al., 2005; Grewe et al., 2017; Gussin et al., 2007b; Nelson et al., 1997; Ratnam and Nelson, 2000). We used a wide range of response parameters to validate our model, including firing rate, coefficient of variation, response sensitivity to random amplitude modulations, response sensitivity and time course to steps, and responses to beat stimuli (see Supplementary Figure S1). This prototypical neuron, with response properties matched to the average experimental values, served as our original seed; the set of parameter values is given in Supplementary Information Table S1.

We then used this original seed to create a heterogeneous population that replicates the range of response properties observed experimentally through an iterative process of diversifying the population and constraining the response properties. A heterogeneous array of parameter sets was created (8,000 sets) from the original seed values by randomly varying slightly several parameters: *t_R_, R_m_, τ_m_*, *V_T_, A_σ_, τ_α_, Δ_α_,* and *β*. From this array, 12 seeds were selected by choosing sets of parameters that, again, best replicate the average response properties. These sets were further diversified randomly, and from these new arrays, 26 seeds were chosen by selecting neurons that replicate the average coding properties but that span the range of spontaneous firing rates measured experimentally (Chacron et al., 2001; Gussin et al., 2007; Ratnam and Nelson, 2000). From these 26 seeds, the parameter sets were further diversified, and we retained 9,200 sets of parameters by rejecting sets for which the response properties did not fit in the range of response properties determined experimentally. Therefore, our pool of neurons replicates the average and range of response properties measured experimentally. From this pool, five equivalent but different populations of 8,195 receptors were created by randomly assigning one of the 9,200 neurons to each receptor location. All data shown in the results reflect averages across these five populations.

### Decoding analysis

Population responses to different stimuli were compared with our decoding analysis to determine how accurately signals could be detected or discriminated, considering the differences in response patterns for these stimuli. The framework used for our decoder has been described and thoroughly validated in previous publications (Marsat et al., 2023; Allen and Marsat, 2018; Allen and Marsat, 2019; Allen et al., 2021; Marsat and Maler, 2010). It was shown that this analysis measure is directly correlated with the information content of the responses about the stimuli. We describe the method briefly here. Importantly, the only difference with the established measure described in Marsat et al. (2023) is that we do not use the detailed time course of each neural response (i.e., the full spike trains) but use the peak-to-though firing rate to quantify the response of the neuron. To calculate the peak-to-trough firing rate, the binarized spike trains are smooth with a sliding square window of 16.67 ms (half a beat cycle) to obtain an instantaneous firing rate. For each beat cycle, the difference between the maximum of the instantaneous firing rate and the minimum gives us our measure of peak-to-through firing rate. These peak-to-trough measures can be averaged over several cycles of the beat, or we can use the values for single beat cycles as specified in the different Results section. When no beat stimulus is provided, the analysis is unchanged, and the peak-to-trough is calculated over each consecutive 33.34 ms segment of response (i.e., a 30 Hz period).

The analysis compares pairs of responses: responses to stimuli from two different azimuths are compared in the angular resolution analysis; while in the detection analysis, the response to the sender fish’s signal is compared to the response when no second fish is present (i.e., baseline firing due to the focal fish’s own signal). For each individual neuron *i* (out of *n* total neurons), the similarity between the probability distributions *P(x)* of responses (the peak-to-trough firing rate *x*) to the two stimuli (*R* and *B*) is calculated based on the area difference *Ω*_(*PR*,*PB*)_ between the two distributions:

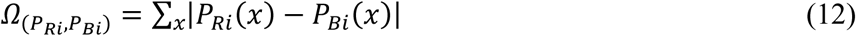

AD values are normalized to 1 across the *n* neurons to obtain a weight W:

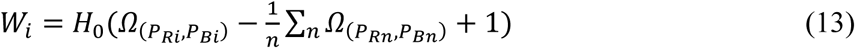

where H_0_ is the Heaviside step function. The peak-to-trough firing rates for each neuron are multiplied with these weights before being used in our Euclidean distance calculations (see below). As a consequence of this weighting, the neurons that respond very differently to the two stimuli will contribute more to the Euclidean distance between the population responses, and the neurons that respond similarly to the two stimuli will contribute little to the Euclidean distance. Since the sum of the weight for a population is 1, the overall firing rate of the population is unchanged by the weighting procedure.

The Euclidean distance *D* between pairs of weighted responses (*R_a_* and *R_b_*) is calculated between responses to the same stimulus or between responses to the two different stimuli being compared:

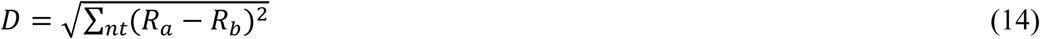

These Euclidean distances are used to determine how well an ideal observer could discriminate between responses to the two stimuli. The distribution of distances between responses to the same stimulus, *D_xx,_* and the distribution of distances between responses to the two different stimuli, *D_xy_*, are used for an ROC analysis. In this analysis, a threshold distance *T* is varied. For each threshold, the probability of non-discrimination (*P_D_*) is calculated as the sum of *P(D_xy_>T),* and the probability of false discrimination *(P_F_)* is calculated as the sum of *P(D_xx_>T)*. The error rate *E* is taken as the minimum error across threshold according to:

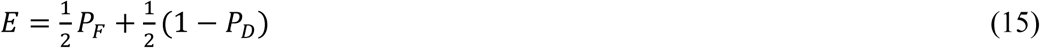

This error rate is used in the various parts of the result as specified therein, and we consider that reliable detection or discrimination happens when the error rate is below 0.05.

## RESULTS

Since ghost knifefish can detect and localize each other based on the electric signals they continuously emit, both fish are thus senders and receivers at the same time. To simplify the description of our results, we describe the fish for which we describe the electric image (EI) and sensory responses as the focal fish. The “other” fish that needs to be detected and localized by the focal fish is designated as the sender fish. A number of studies have characterized the EI that one fish causes on another fish’s body (Kelly et al., 2008; Pedraja et al., 2016). The results presented in these papers are not always easily related to electrophysiological studies because many experiments on the responses of sensory neurons in this system calibrate the signals as a relative contrast in the measured voltage. For example, many studies present the response properties to a conspecific signal of 5-10% contrast (Fotowat et al., 2013; Metzen et al., 2018). Since our goal is to use the estimate of signal strength as an input to neuron models that match their known response properties, we quantify the EI strength as a relative contrast in the transdermal potential difference. If the signal from the sender is as strong as the focal fish’s own baseline signal at a given point on their body, the signal strength will be 100%, and the EOD will vary from close to 0 mV (at the trough of the beat) to twice the baseline EOD strength (at the peak of the beat cycle).

Expressing the signal as a contrast highlights the challenges that the fish encounters when interacting with a conspecific due to the rapid decay of signal strength with distance. Figure 1 displays the signal strength during a common scenario: a sender fish approaching and then moving away from the focal fish. When the fish are separated by only a few centimeters (position 2), the EI has a strong gradient that goes from a strong signal (25% contrast) on portions of the body closest to the sender to a very weak signal close to 0% contrast. For convenience, we will refer to the region of the electric image with the strongest signal as the hot spot. While this hotspot is well-defined when the sender is close by, the EI is much more diffuse when the sender is further away, and there is a weaker gradient between the hotspot and the areas with a weaker signal. For example, in Fig 1B, the difference between the hot spot near the head (position 1) or tail (position 3) is less than 1% stronger than the portions with the weakest signal. The positions 1 and 3 depicted here correspond to a distance (30 cm) at which we know fish can detect each other and display active interactions (Henninger et al., 2018; Zupanc and Maler, 1993; Zupanc et al., 2006). The known extreme sensitivity of this system is thus highlighted here, as detecting the sender 30 cm away involves detecting a 1% modulation, and localizing this signal requires deciphering a gradient in the signal of less than 1% across the body surface.

**Figure 1:**
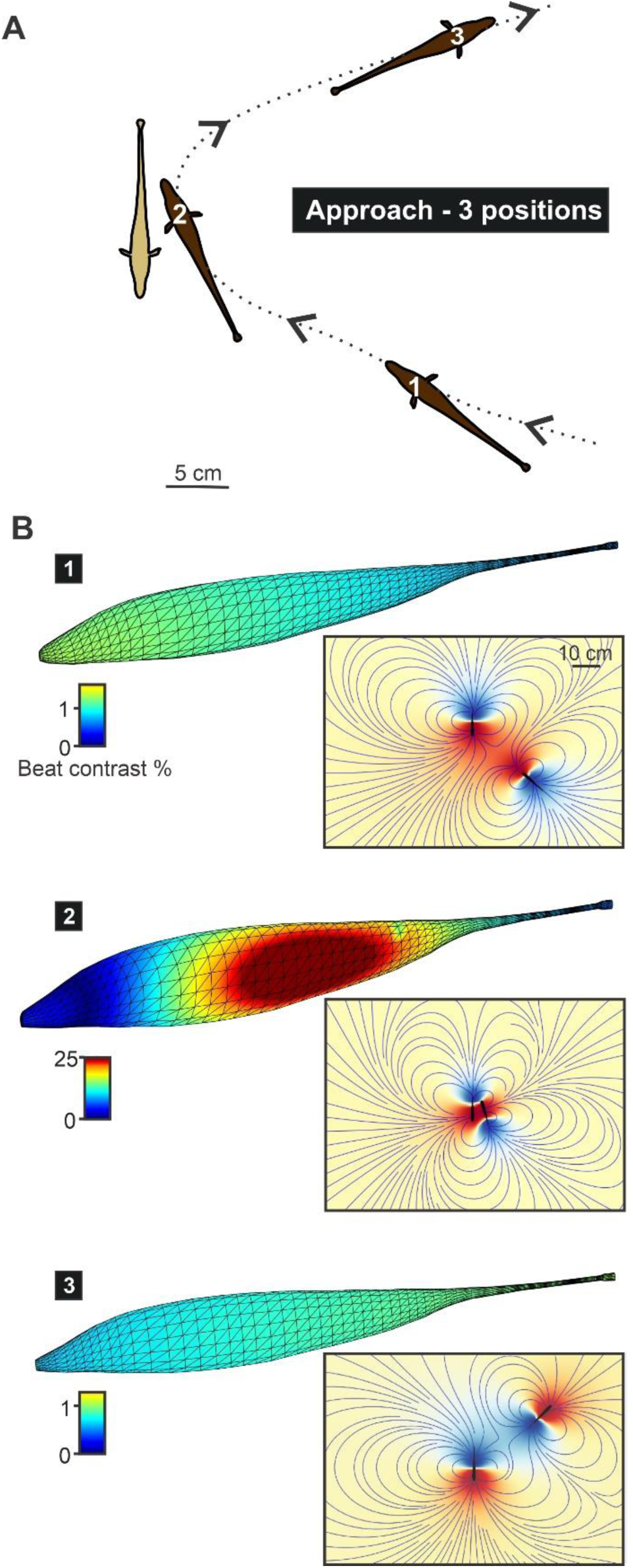
Model of the electric image during social interactions. The model takes into account the relative position of the two fish to estimate the strength of the EI caused by one fish (designated as “sender”) onto the body of the other (“focal” fish). **A.** Three relative positions are illustrated here, representing different phases of a fish approaching the focal fish. **B.** The EI is quantified based on the strength of the transdermal voltage at the peak of the beat AM caused by the interaction of the fish’s EODs. The strength of this AM is then normalized to the baseline EOD strength (i.e., EOD amplitude when only the focal fish is present) to obtain an EI strength expressed as percent contrast. 0% contrast indicates that the signal from the sender fish has no impact on the receiver, and 100% contrast indicates that the sender’s signal is as strong as the focal fish’s own EOD. We can see that for the two more distant positions, the signal strength is weak (<1%), and there are only minute differences in signal strength across the receiver’s body. When the sender is very close, however, a salient “hot spot” has a much stronger EI strength than other portions of the body (note the different color scales for each image). We show, as an inset on the right, the electric field with current lines (grey lines) and the iso-potential lines, depicted as a color gradient (red or blue depending on polarity), for the near-field range. The perspective in this inset is from the top as in A), while the EI illustrations in B) present a perspective of the side of the focal fish being approached by the sender.

The EI stimulates an array of receptors covering the fish’s skin, and the spatial structure of this sensory array will dictate how the spatial structure of the signal is captured. Receptor density for different portions of the fish’s body has been characterized in a closely related species (Carr et al., 1982), but we needed a more detailed quantification of variations in receptor density across the fish’s body (Fig 2A). Our data confirm the general organization of receptor density: a region of high density on the snout and head, of medium density on the dorso-rostral portion of the trunk that decreases both ventrally and caudally (Fig 2B). Based on this spatially precise empirical data, we incorporated a population of p-unit receptor locations on the 3D mesh model used for EI calculations that varies smoothly in density as a function of rostro-caudal and dorso-ventral position (Fig 2C). In the rest of this study, we will use an “average” fish that measures 14 cm long and includes ∼8,195 receptor locations.

**Figure 2:**
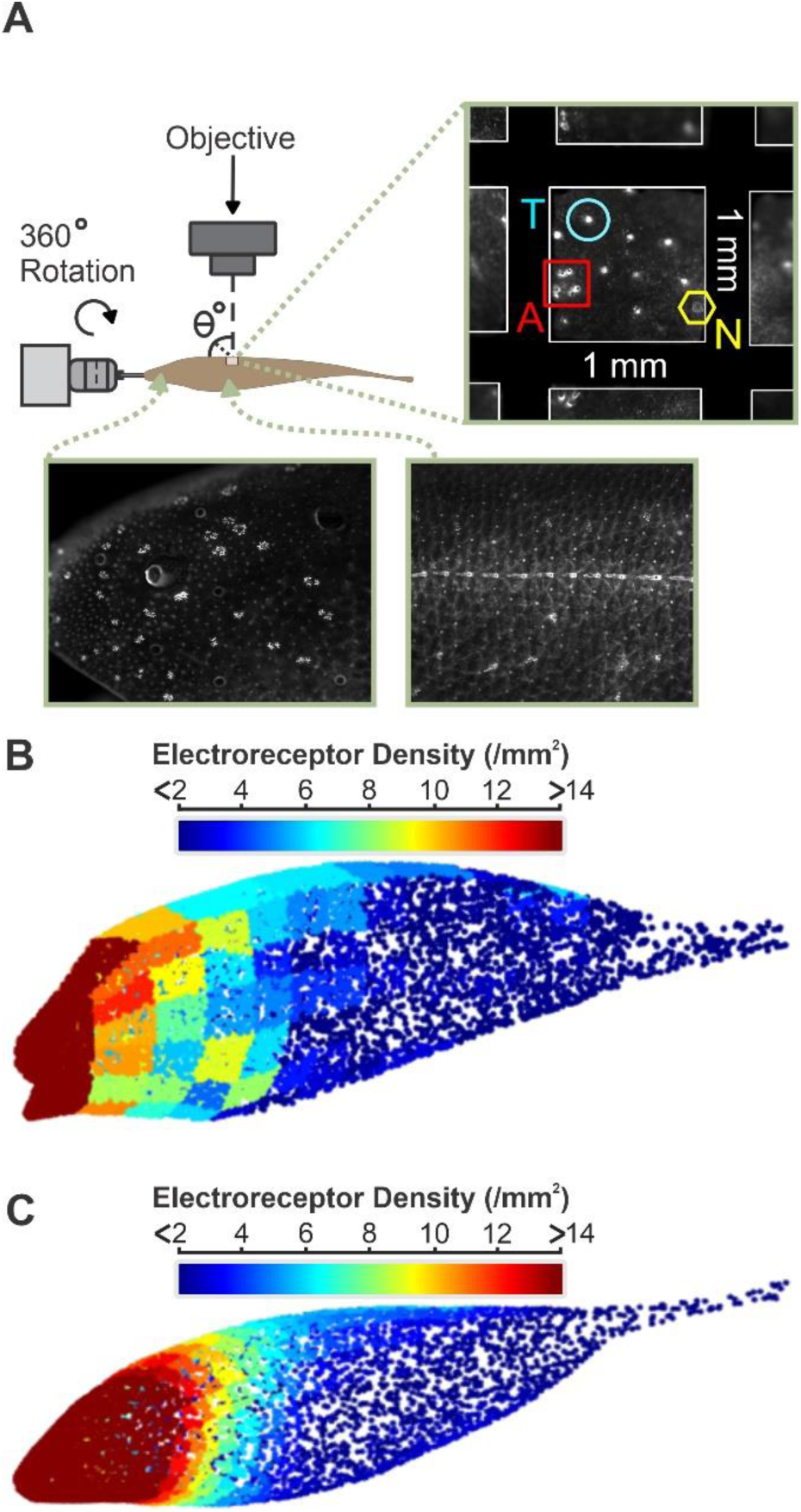
Spatial structure of the receptor array. **A.** Receptor density across the body of the fish was determined experimentally. Eosin Y stained specimens were examined, and cutaneous receptors of different types were identified (e.g., Neuromast labeled N, ampullary labeled A, or tuberous receptors labeled T in the inset on the top right). The number of tuberous receptors in 1mm^2^ areas was averaged across samples for each face of a coarse 3D mesh model of the fish (**B**). Receptor density as a function of body position was then smoothed by fitting a 5^th^-degree polynomial and mapped onto the fine mesh model used in the EI model (**C**). For each face of this 3D mesh, random receptor positions were selected according to the receptor density attributed to each face. The receptor positions that we generated are marked by the dots in B) and C), and their color reflects the density of receptors for the corresponding position.

We calculated the stimulus intensity for each receptor location for the various iterations of the EI calculation; the result for the three positions illustrated in Fig 1 is shown in Fig 3A (note that for this figure, the blue-red color scale covers the contrast range for each position). We wanted to quantify how the strength of the EI changes with the position of the sender relative to the focal fish. To do so, we ran the model for 864 relative positions, where the two fish are at 12 distances that vary from nearly touching to 75 cm apart, and for 72 different azimuths around the focal fish while always having the sender fish’s heading towards the focal fish. To estimate the maximal strength of the signal for each location, we used the strength of the signal at the center of the hot spot on the focal fish. We plot this value as a color scale at the position of the middle of the electric organ of the sender fish (Fig 3B). We can see that the signal strength decreases sharply with distance, as expected. The portions of the figure with contrast above 20% (yellow-white shades) represent positions where the fish are nearly touching (with the sender fish head-on). The decrease in signal strength with distance follows the expected power law, such that contrast drops to 10% by 10-15 cm and is only a few percent when the sender is 20-30 cm away (Fig. 3C).

**Figure 3:**
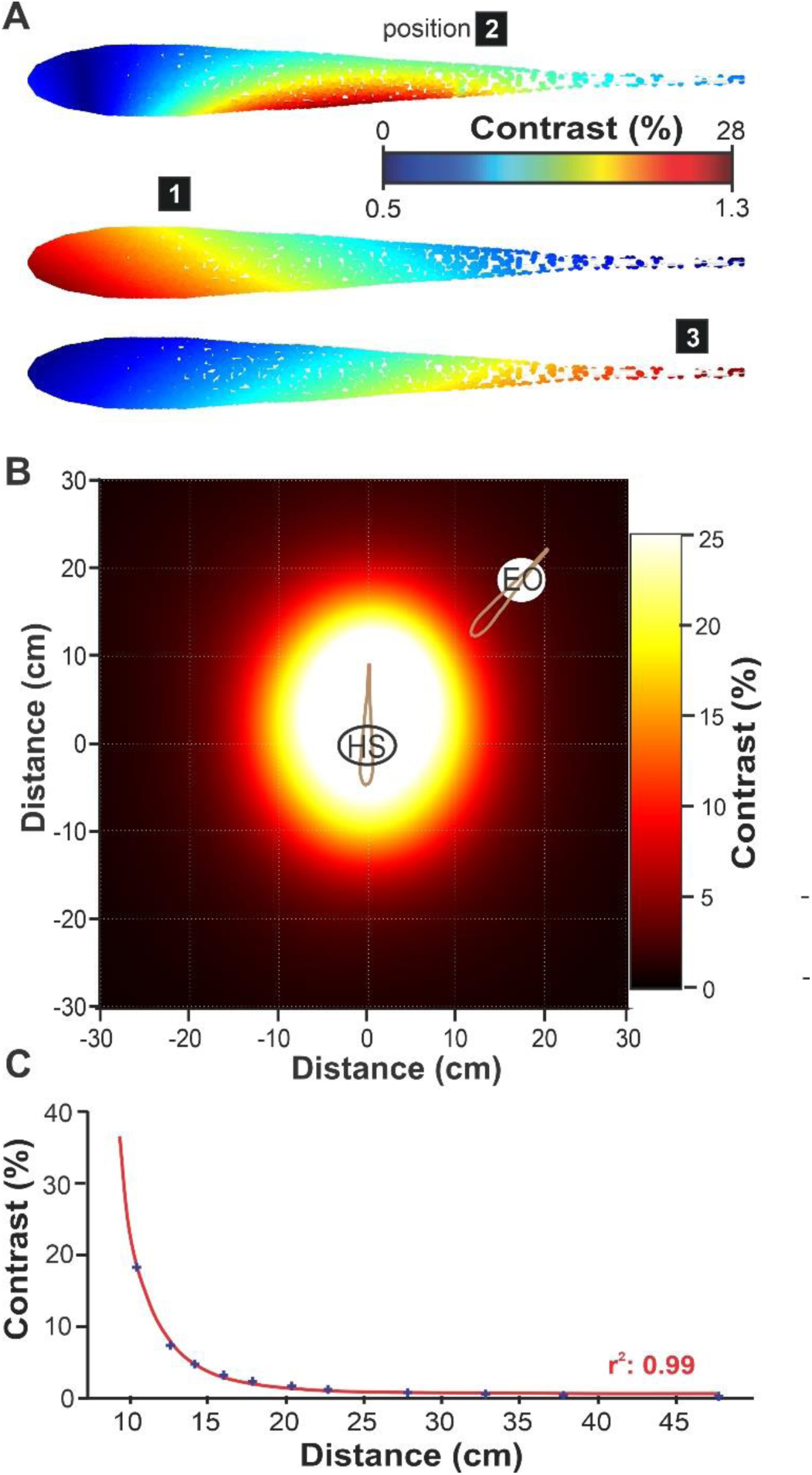
EI strength as a function of the relative position of the two fish. **A.** The strength of the EI from the sender was calculated for each receptor position. It is displayed here for the three relative fish positions illustrated in Figure 1 (note the difference in color scale for positions two vs. one and 3). The focal fish are presented here from a top perspective. **B.** The strength of the electric image is characterized as a function of distance and azimuth. The EI for 864 relative positions (12 distances x 72 azimuths) was calculated, and the strength of the signal in the “hot spot” (HS; taken as the average of the 5% most strongly activated receptors) is depicted by the color scale. The position of the sender fish is based on the center of the EO, and the data points have been repositioned to reflect the distance between the center of the HS on the focal fish and the EO of the sender rather than the center of the focal fish. The EI strength values (contrast %) have been interpolated between data points. **C.** Average contrast values in the HS across azimuth and for fish displayed as a function of the distance between the hot spot and EO centers. The relationship follows a cubic power law (red best-fit curve: *c=1.6·10^4^·x^−3^*).

The EI model helps us quantify the cues that the sensory system can use to detect and localize the source of these signals. To better understand the accuracy with which detection and localization could occur, the strength of these signals must be compared with the noise that the sensory system experiences. We therefore use the deterministic model of EI signal strength described above as an input to a population of model neurons that includes realistic noise. We based our receptor model on established parameters of leaky-integrate-and-fire, which include adaptation, noise, and a refractory period, to replicate the response properties of p-units (Benda et al., 2005; Chacron et al., 2005; Grewe et al., 2017; Gussin et al., 2007; Nelson et al., 1997). Based on this prototype p-unit model, with properties matched to the average characteristics of p-units, we diversified the model parameters to create a heterogeneous population matching the range of properties observed experimentally (see Methods for details). For the model to provide a biologically reasonable estimate of how accurately this population encodes spatial cues, it is vital to calibrate the sensitivity, heterogeneity, and noisiness of the responses to reflect the measured properties of the neurons. Our calibration involved matching the model responses to published values for a range of stimulus types and analysis methods (see Methods and Supplementary Figure S1). The population is then stimulated with conspecific signals (beat stimuli) at intensity levels that are dictated by the EI image for various relative fish positions. Figure 4 shows the response pattern of the population for two of the positions used in Figures 1 and 3. Response strength is quantified as the difference between the peak and trough of the instantaneous firing rate during each stimulus cycle (here normalized relative to spontaneous activity). We can observe differences in response strength due to the spatial contrast in the EI, as well as differences across receptors resulting from heterogeneity that makes some neurons more sensitive.

**Figure 4:**
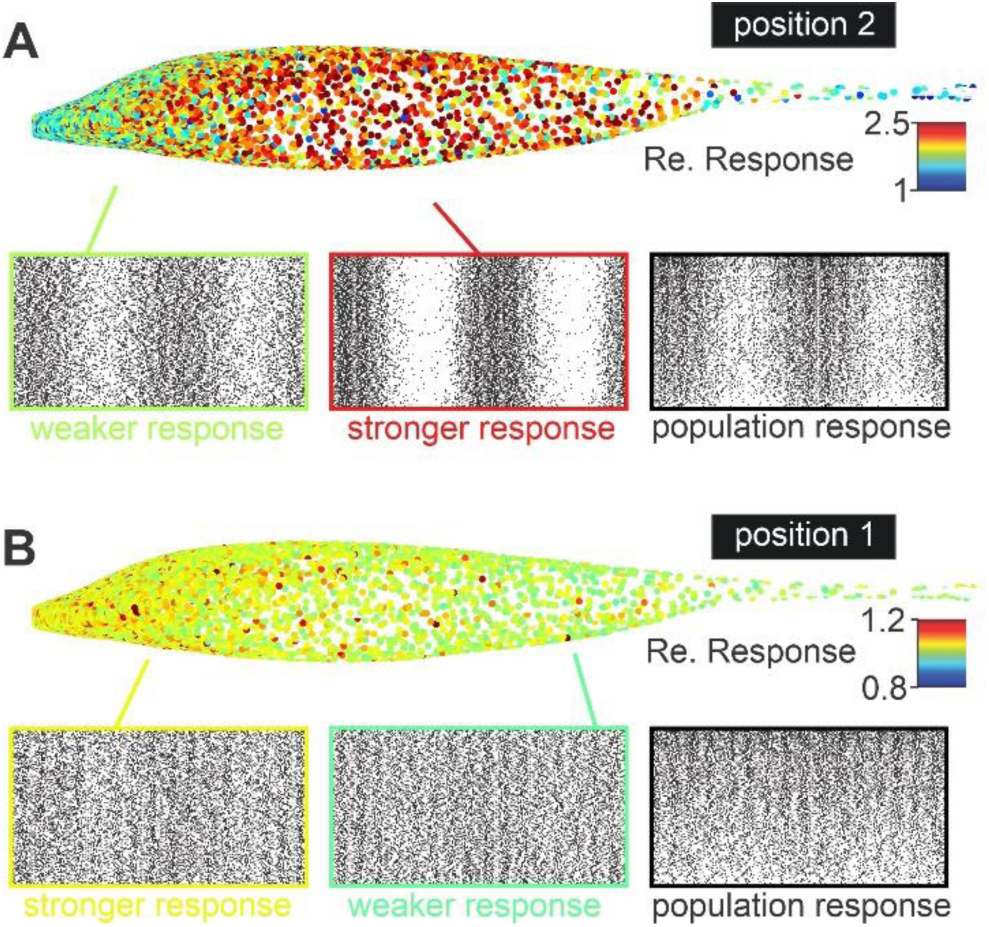
Heterogeneous population response in modeled receptors. **A.** LIF models with heterogeneous response properties were stimulated with realistic inputs that match the spatio-temporal structure of conspecific signals. In the color map (top), each receptor’s response is quantified as the peak-to-trough firing rate modulation and normalized relative to spontaneous modulations in firing rate occurring when no second fish is present (i.e., a relative response of 1 reflects modulations in firing rate similar to spontaneous activity). The two raster plot insets on the left show the response patterns of two individual neurons from portions of the fish’s body that are more or less strongly stimulated. The raster plot on the right shows the population response, where we display a stack of 800 randomly selected receptor responses, ordered from weakest peak-to-trough responses (bottom) to strongest responses (top). **B.** We display the same elements as in panel A, but the position of the sender fish is more distant (position 1 of Figure 1) compared to the nearby relative position used in A. (position 2 from Figure 1).

Firing rate modulation across one stimulus cycle replicates the experimentally measured sensitivity (Fotowat et al., 2013; Henninger et al., 2018; Pedraja et al., 2014; Pedraja et al., 2016; Zupanc et al., 2006) The strength of the electric image quickly decreases with distances and reaches levels below 1% contrast at ranges where the fish still detect and interact with each other (e.g., 30 cm). Our model replicates the fact that for these weak signals, the response modulation is barely above variations in firing rate that naturally occur in the absence of a second fish (Fig 5). Figure 5C shows that, although the responses for nearby fish (14 cm and 22.5 cm in this figure) are visibly different from the spontaneous response, signals from more distant fish (40 cm, 75 cm in this figure) cause much more subtle differences in response strength. This is true when looking at the distribution of responses for cycles of the beat, for responses averaged across time (grey or color portion of the distribution plots), or overall responses averaged across time and neurons (white lines).

**Figure 5:**
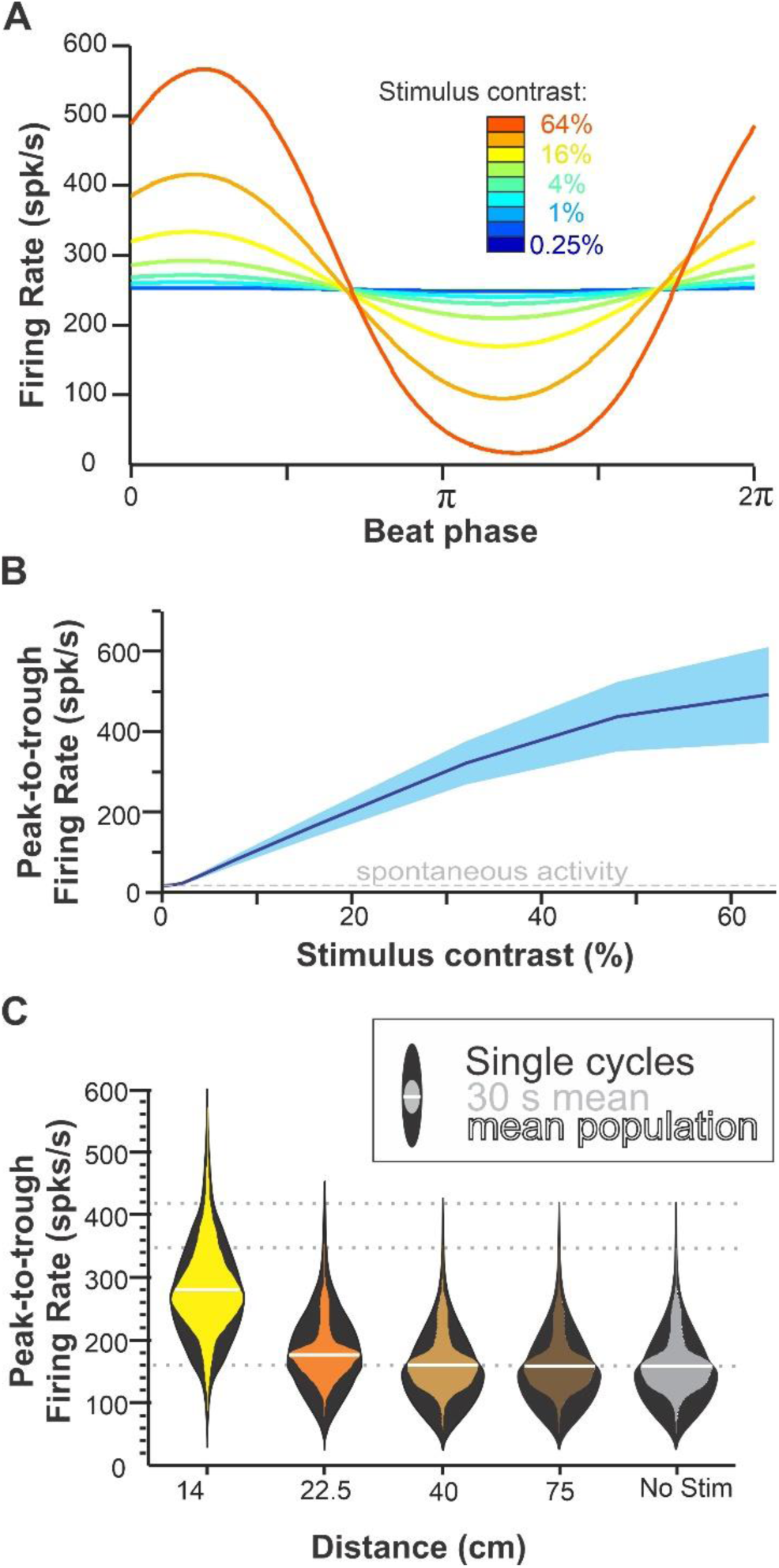
Sensitivity of the receptor model replicates responses to beat stimuli. **A.** Mean firing rate (averaged across all receptors) during a single cycle of a 30 Hz AM beat stimulus for different contrast intensities. The model produced modulations in firing of just a few spikes/s peak-to-through for the weakest intensities and of a few hundred spikes/s for the stronger intensities. **B.** F-I curves display the strength of the response as a function of the stimulus intensity (mean across the population ± s.d.). Average peak-to-through firing rate quantified in a similar way; spontaneous activity (no sender fish) is displayed (grey dashed line), and we can see that for the weakest stimuli the average response is similar to spontaneous activity. Response sensitivity has been calibrated to match published data (Bastian, 1981; Nelson et al., 1997; see also Methods and Supplementary Figure S1). **C.** The distribution of response strength across the population of receptors is displayed for sender fish positioned at different distances (azimuth 90°). The outer distributions (black violin plots) show the variability of peak-to-through firing rate on single cycles of the beat for single neurons. The inner distributions (colored violin plots) show the variability across neurons; however, for each neuron, the peak-to-trough firing rate is averaged across the entire stimulus (900 cycles, 30 s). The mean of this distribution (i.e., also averaged across neurons) is displayed as a white line. Dotted grid lines (in grey) are provided in the background to help notice minute differences in the distributions. The fact that only subtle differences between the distributions for stimuli at distances of 40 cm or more compared to responses in the absence of second fish highlights the difficulty in detecting these distant signals.

The sensitivity with which a conspecific signal would be detected will depend on the coding and decoding mechanisms implemented by the nervous system. It is beyond the scope of this article to explore all possible coding and decoding algorithms that could contribute to the system’s sensitivity, but as a first step towards understanding the system’s sensitivity, we aim to provide a lower bound on accuracy. To do so, we only consider peak-to-trough firing rate to quantify response strength as it is the most salient aspect of the response that varies with stimulus strength. The responses of individual receptors will be combined at higher levels of the nervous system, and this can help average out noise in the responses. However, the way these responses are combined would require various assumptions matched to the architecture of high brain areas, which would lead to a complex modeling effort and speculation about the decoding procedure. For this reason, and to remain within a “lower-bound” perspective, we map each population response in Euclidean space, where each dimension represents the response strength of a neuron (i.e., response strength is not averaged across neurons). Reliable detection would occur if the strength of the population response is markedly different from the response when no signal is present; in other words, when the stimulus-response is far from the baseline-response in this Euclidean space representation. Our analysis, therefore, uses a weighted Euclidean distance followed by a ROC analysis to quantify the reliability with which stimulus responses can be differentiated from baseline. This analysis could be conducted on “instantaneous responses,” considering the response of the population (peak-to-trough for each receptor) for a single cycle of the stimulus, which would give us a response accuracy (i.e., probability of error) for each fish position. Behavioral responses typically occur after attending the signal for a certain period of time, and more accurate detection occurs after several seconds of the stimulus than after the first cycle. We therefore integrate the response of each neuron across time (averaging the peak-to-trough across several beat cycles) to estimate how detection accuracy changes as information is accumulated over time. We consider that accurate detection occurs when less than 5% detection error would occur and plot in Figure 6 the amount of time that accurate detection would require. Our analysis shows that accurate detection could occur within a single cycle of the stimulus for fish that are within 20 cm of the focal fish (Fig 6A, 6C). Detection accuracy decreases with distance, and thus, it takes more time to detect the stimulus reliably. As a result, a fish 60-70 cm away would require integrating the signal for several seconds to be able to tell reliably that another fish is present.

**Figure 6:**
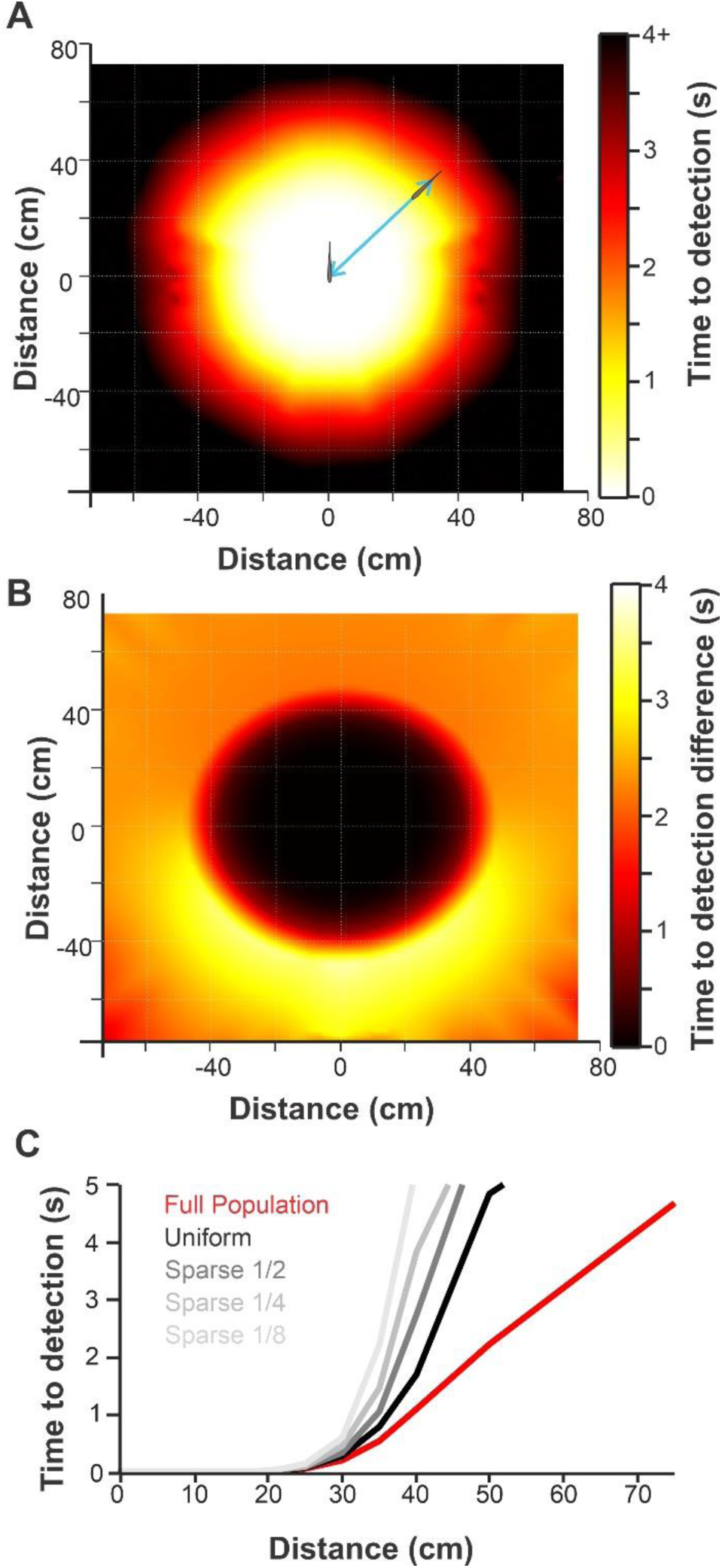
Estimates of detection sensitivity as a function of relative fish position and the influence of receptor density distribution. **A.** Detection sensitivity is quantified as the time it would take to reliably detect the presence of the sender’s signal and is displayed as a function of the relative position of the sender. For each position (12 distances x 72 azimuth), the peak-to-trough responses of the population are compared to spontaneous responses. As peak-to-trough is averaged across cycles of the beat (increased time to detection), detection becomes more reliable because noise is averaged out. We display the stimulation time required for our decoder to reach reliable detection (<5% error; see Methods for more details). Each position is defined by the distance and azimuth between the center of the receptor positions in the focal fish and the center of the EO in the sender fish. Values for each data point are then interpolated across space to obtain the smoothly varying color plots. **B.** Decrease in detection sensitivity (increased time to detection) caused by reducing the density of receptors in dense areas (i.e., making it uniformly low like the density in the caudal portion of the body). The decrease in density makes uniformly no difference nearby (black region). It uniformly increases the time to detection when the sender is very distant (distant black area in panel A). It has a most pro nounced effect at mid-distance (40-50 cm) in the frontal azimuth (yellow area) where time to detection increases from 1-2 s to 2-6 s (see also Supplementary Figure S3) showing that the rostral high density of receptors can increase detection sensitivity in the frontal quadrant. C. The overall detection sensitivity (time to detection averaged across azimuth) decreased markedly at distances above 30 cm when receptor density was made uniformly low (black line compared to red line) and decreased further when receptor density was made even

Detection accuracy is undoubtedly influenced by the fact that thousands of receptors contribute to transmitting this information. The distribution of these receptors is not uniform across the fish’s body. In particular, the rostral end of the fish (head region) has a density of receptors 5-10 times higher than the caudal end (tail region). We hypothesize that this foveal organization supports an enhanced sensitivity in the frontal quadrant relative to the fish. Alternatively, the increased density in receptors plays an important role in localizing small objects, like prey, with higher resolution but does not enhance the perception of conspecifics in specific regions of space. This is a plausible alternative because conspecific signals cause a more diffuse EI that stimulates a majority of the receptors over the receiver’s body. To test our hypothesis, we reduced the density of receptors to make it uniform over the entire body and equal to the low density found at the caudal end of the fish (Supplementary Figure S2). Our resulting population has 2,770 receptors compared to 8,195 for the full population. The difference in detection sensitivity is displayed in Figure 6B and reveals the enhancement in sensitivity afforded by the increased density in the rostral portion of the body. The difference is negligible when the sender is very close because accuracy is high, and a decrease in the number of receptors encoding the stimulus is not sufficient to affect performance. For very distant signals (e.g., 75 cm), the lower rostral density causes a decrease in sensitivity that is fairly uniform across azimuth (see also Supplementary Figure S3). This can be explained by the fact that distant signals cause diffuse EI patterns that vary by less than 1% across the body (e.g., see Figure 1). As a result, the head region is stimulated at a level nearly identical to the tail region, even if the sender fish is located behind the focal fish. There is, however, a clear difference as a function of azimuth for medium distances (35-50 cm; Figure 6B and Supplementary Figure S3). The higher density of receptors on the head provides a clear advantage in detection accuracy in the frontal quadrants (i.e., it requires less time until accurate detection occurs). We further verified the idea that having a larger population of receptors allows the population to encode the presence of the stimulus more reliably by decreasing the density of our uniform population of receptors by 2-, 4-, and 8-fold (resulting in population sizes of 1,385, 693, and 347 receptors, respectively). Our analysis confirms that a higher density of receptors allows more accurate detection for distances where the signal is faint (Figure 6C).

We next inquire about the spatial information about the angular position of the sender fish relative to the focal fish: its azimuth (in front=0°; behind= 180°). To do so, we compare the responses to stimuli at various positions and quantify how reliably these responses could be discriminated based on the same weighted Euclidean distance analysis used above. Error probability will thus depend on two factors: the angular separation between the two stimuli and the duration of the stimulus evaluated, whereas in our previous detection analysis, the latter was the only factor considered. To display our results conveniently, we chose to keep the duration of the stimulus integration at 1 second. This value is a compromise between expecting accurate angular discrimination instantaneously (∼1 cycle) and expecting the sender fish to remain in a relatively fixed position for seconds in order for the angular position to be accurately estimated. Using this 1s integration time, we determined the angle that needed to separate two stimuli to lead to reliable discrimination (<5% error) and refer to this value as angular resolution (Figure 7). Our results show that angular resolution is accurate to a few degrees (<5°) when the sender is within 10-20 cm and decreases sharply between 20 and 40 cm such that beyond 40 cm, azimuth could not be reliably determined (angular resolution>180°). If we repeat the analysis using different integration times (Fig 7B), as expected, the sharp decrease in angular resolutions moves from being for positions 20 cm away when a single cycle of the beat is considered to being 40-50 cm away when 3.3s of the stimulus is being averaged. Our analysis suggests that for the most distant positions tested (75 cm), the angular position could not be resolved, even when the stimulus was integrated for several seconds. Together with the results of Figure 6, these findings suggest that for more distant signals, only detection would occur, and the position of the sender would not be accurately determined without relying on additional behavioral or neural mechanisms (see Discussion).

**Figure 7:**
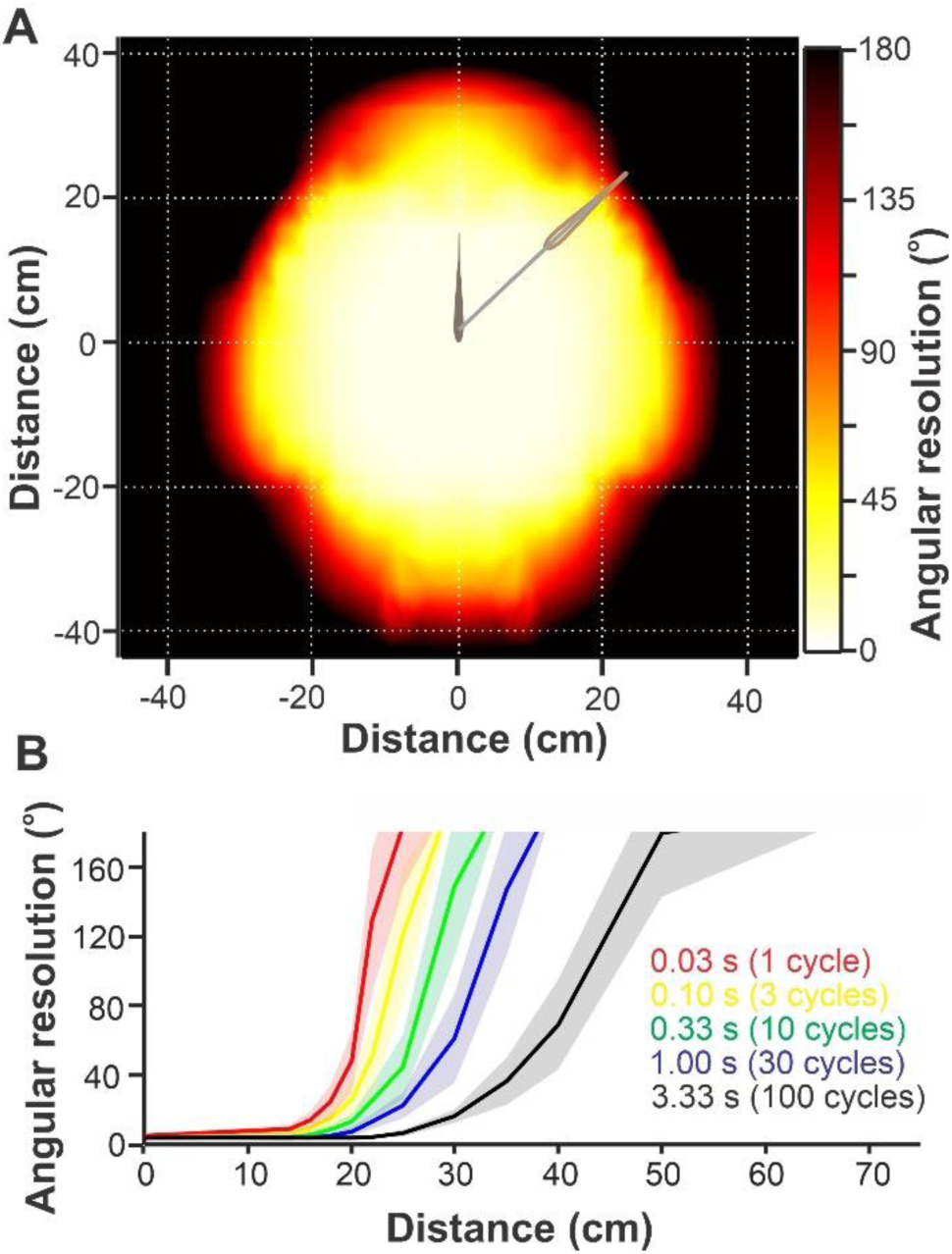
Estimates of angular resolution supported by the receptors’ response accuracy. **A**. Angular resolution estimated as a function of distance and azimuth of the sender. The analysis is based on the average peak-to-trough firing rate (30 cycles average, i.e., 1 s) of each receptor for a given stimulus location. Population responses for a sender at a given test azimuth were compared by our decoder to responses for a sender at various angular displacements (clock-wise or counter-clock-wise; distance kept fixed). The smallest angle that allows 95% accurate discrimination between the responses was taken as the angular resolution. Failure to discriminate responses for stimuli locations 180° apart (black regions on the graph) indicates an inability to reliably localize the azimuth of the sender’s location. Data points for 72 test azimuths and 12 distances (i.e., 864 positions) were generated, taking the center of receptors location for the focal fish and the center of the EO for the sender fish as position values. Data values are then interpolated to obtain the smoothly varying color plot. **B.** The angular resolution (mean ± s.d. across azimuth) can be calculated based on peak-to-trough responses of the receptors to a single cycle of the stimulus or on the peak-to-trough averaged across several cycles. Averaging the response across cycles corresponds to a decoder that integrates the response across time to get a more accurate estimate of response strength. Consequently, the angular resolution improves as the decoder integrates over more time; however, in all cases, we still observe a sharp change in resolution from very accurate (e.g., at distances below 20 cm) to very poor (e.g., above 50 cm).

The angular resolution does not appear to be equally good for a given distance as a function of the azimuth of the sender (e.g., front vs. side vs. back). This can be seen in Figure 7A, where the color patterns around the focal fish are not perfectly circular. In particular, worse angular resolution occurs at a shorter distance in the back quadrant compared to the front. We suspected that the way we equalized distance across angles could cause a bias. The center of the focal fish is taken as the average receptor position, and the center of the sender as the middle between the two emitting poles that constitute the EOD source. Center-to-center distance and azimuth were used to set our relative positions. In this arrangement, a fish at 130° (in the back quadrant, see fish illustrated in Fig 7A) rotating 5° would not only change azimuth but also come closer to the focal fish if we consider the two closest points on each fish’s bodies. Note that this is not an issue in our detection analysis since our accuracy measure does not depend on the comparisons between two stimuli locations but only on the absolute location of a single stimulus. We therefore repeated our angular resolution analysis by comparing angular resolutions where the distance between the rostral tip of the sender and the closest point on the body of the receiver is kept constant (the dots in Fig 8A show the positions of the rostral tip of the sender used in our analysis). We found that angular resolution is better in the frontal quadrants than at the back (Fig 8A, 8C). Surprisingly, it is not best directly in front of the focal fish (0°) but rather on the side (90°). Since we expected the high density of receptors on the head of the fish to help enhance localization accuracy, we hypothesize that angular resolution is particularly good as the edge of the hotspot caused by the sender sweeps across the region of high receptor density (which could correspond to sender positions around ∼90°). To test the contribution of higher receptor density towards the head, we repeated the analysis using the “uniform” receptor population from the previous section, which has an equally low receptor density across the whole body. This uniform population of receptors did not perform much worse than our full population, and in particular, there is no striking difference directly in front (0°) or the side (90°) of the fish where resolution is best. Azimuth determination was not possible (angular resolution > 180°) when the fish were two body lengths apart (28 cm), but at one body length, the higher density of receptors provided a small advantage. Specifically, there are only a few spots at ∼45° and ∼125° where the higher receptor density on the rostral portion of the body helps to enhance angular resolution (Fig 8B, 8C). We confirm that decreasing the population density further decreases angular resolution, and this is particularly evident for more distant stimuli (one body length), where localization is still possible but the angular resolution is not great (Fig. 8D).

**Figure 8:**
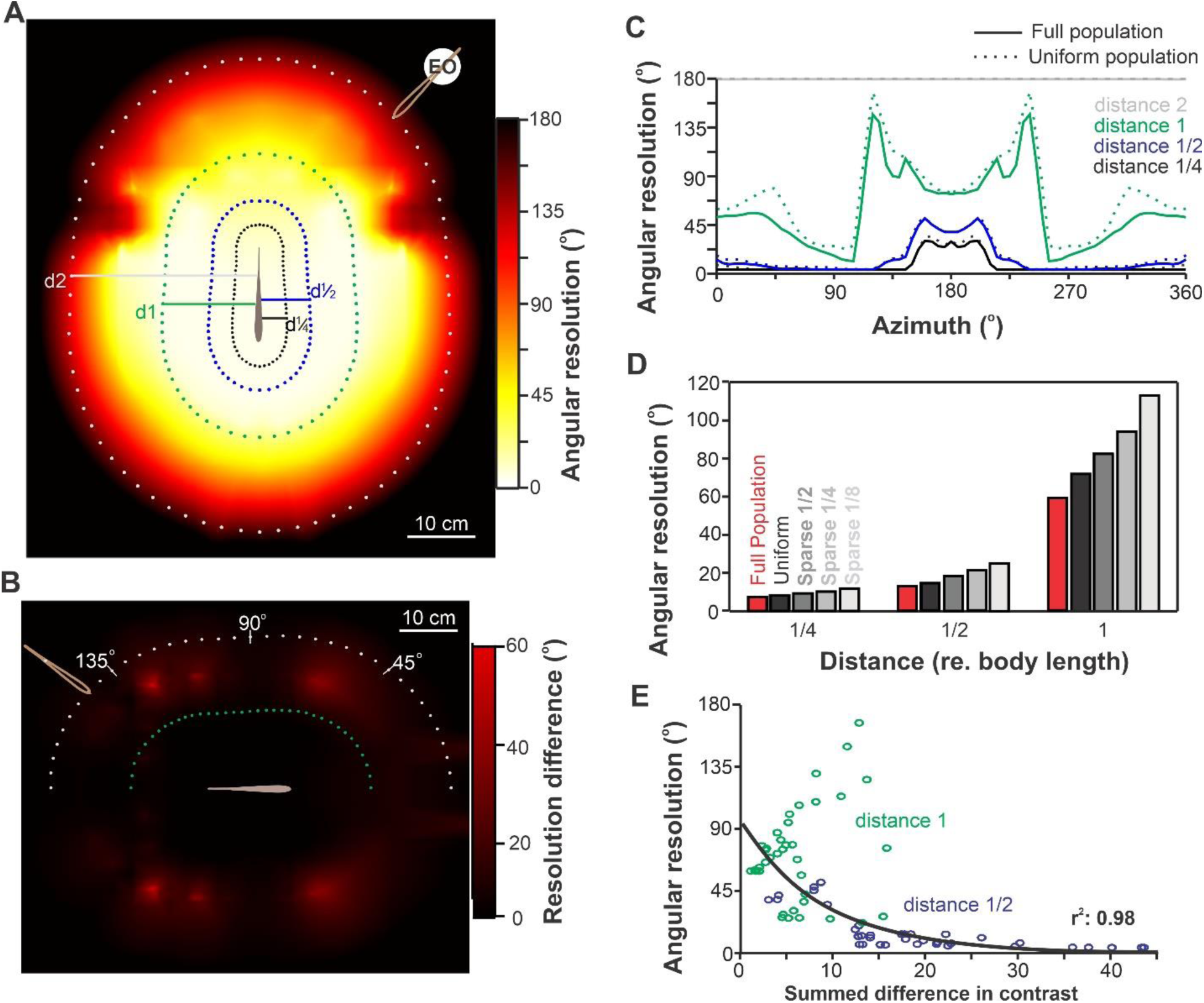
Receptor density and angular resolution as a function of azimuth. **A.** Angular resolution for a sender fish at various positions around the focal fish was calculated as in Figure 7, but the way distance is normalized across angles is different. We still compare, with our decoder, positions at different angles while keeping the distance equal. Here, distance is set by the distance between the rostral tip of the sender and the closest point on the surface of the focal fish. The fish are thus separated by a fixed gap set to 1/4, 1/2, 1, or 2 body lengths (i.e., 3.5, 7, 14, and 28 cm, respectively). In Figure 7, the gap between the fish was not consistent across azimuth since the distance was set between the center of receptor locations in the focal fish and the center of the EO in the sender. For example, the rostral tip of the sender was closer to the skin of the receiver when it was at 180° than when it was at 0°. The positions of the rostral tip of the sender, for the 72 angles and four distances, are marked with dots on this figure. Datapoints for various positions are still mapped on the color graph at the position of the middle of the EOD of the sender, and the color gradient interpolated between datapoints. **B.** Decreasing the density of receptors in dense areas (i.e., making receptor density uniformly as low as density in the caudal portion of the body) causes decreases in angular resolution. This resolution decrease is strongest in the red area of the graph. We note that this decrease is relatively limited and not concentrated in the frontal quadrant. **C.** Angular resolution as a function of azimuth, distance, and receptor density. This is the same data used to generate Panels A and B but presented here in 2D. The change in resolution as a function of azimuth is clearly visible even when receptor density is uniform across the body **D.** Angular resolution averaged across azimuths for different receptor density patterns. The full population is compared to a uniform lower-density population or to populations made even sparser (1/2, 1/4, or 1/8 of the uniform population). Decreasing the receptor density decreases the angular resolution. **E.** Changes in EI for small angular displacements (averaged across 5°, 10°, and 15° displacements) are compared across azimuth and related to the angular resolution of the system. For a given displacement, the difference in EI was characterized by integrating the difference in EI strength across the body surface. This EI difference was calculated for various azimuths and at the two distances (1 or 1/2 body length apart) that give medium angular resolutions. We show that the EI difference correlates with angular resolution with an exponential relationship (black best-fit curve, r^2^=0.98).

Our results demonstrate that the angular resolution is relatively poor at the back and is slightly better on the side than at the front. This effect cannot be attributed to the higher density of receptors towards the head. Therefore, we questioned whether this effect could be due to the geometry of the fish’s body and how it interacts with the geometry of the electric field. We hypothesize that a small change in angle on the side will cause a relatively bigger change in the electric image than a similar angle difference at the back. A sender fish at 90° azimuth would have its EI hotspot centered on the flat surface of the side of the focal fish, whereas for frontal (0°) or caudal (180°) azimuth, the EI hotspots are centered on the pointy rostral and caudal ends of the fish. We quantified how much difference in EI a sender at various azimuths would cause on the focal fish and integrated this difference across the body surface. We found that it correlates strongly with the differences in angular resolution that we have estimated for various azimuths and distances (Figure 8E). Our analysis supports the conclusion that differences in angular resolution as a function of azimuth are, in part, due to the geometry of the fish and their electric fields.

## DISCUSSION

By using a model of weakly electric fish EI and carefully normalizing how we calculate the distance between the relevant points on the two fish, we presented a clear quantification of the strength of the EI as a function of distance. Our results suggest that strong signals of more than a few percent beat contrast occur only at distances below 15-20 cm. This finding is in agreement with empirical data and consistent with the fact that when two fish actively interact (e.g., chasing each other and courtship), they are typically in close proximity (Fotowat et al., 2013; Zupanc and Maler, 1993). The relationship we show is quantitatively informative only in a simplified case: a given fish size (and EOD strength) with a fixed heading angle (sender head-on towards focal fish). The strength of the electric image as a function of distance will depend on environmental factors (water conductivity), the individuals interacting (their size and EOD strength), and moment-by-moment changes in the relative heading angle of each fish. All these factors could be taken into account in a more extensive analysis of EI during social interactions, but it is beyond the scope of our paper. It is also important to note that further improvements to the model could provide additional details on the structure and dynamics of the EI, such as replicating the bending of the fish’s body or modeling the EO with more spatio-temporal details rather than being a simple fixed dipole. We note that since the strength of the EI at a given receptor location depends on distance and relative heading angles, the strength of the beat AM (its envelope) cannot serve as a reliable indicator of distance or movement towards/away. Rather, reliable spatial information must take into account the differences in EI strength over the body of the receiver. This is obvious for localizing the azimuth of the target but can also help resolve the distance since a fish close by will cause EIs with sharper contrasts across the body, while distant fish will elicit more uniform EIs.

We used, for the first time in this system, a model of the full population of receptors, replicating their heterogeneous response properties and their spatially realistic input patterns. This allowed us to provide a conservative estimate of the detection range that this sensory input could support. We found that, depending on how the information is extracted, detection would still be possible at 75 cm. This range is comparable to behavioral interactions that report instances where a fish detected and moved towards another, or simply interacted electrically,t ranges above 60 cm (Henninger et al., 2018; Stamper et al., 2012; Yu et al., 2012). Our sensitivity estimates might, in fact, come short of the sensitivity observed behaviorally, but this might be because we intentionally provide a conservative estimate. Our estimate relies on a decoding analysis that does not try to replicate sophisticated decoding procedures that the nervous system could implement. Our analysis uses a “Euclidean distance” perspective to quantify similarity in responses where the response of each neuron is kept as separate dimensions. The nervous system will, in various steps of its pathway, combine neural responses and thereby average out noise. An optimized procedure (e.g., using a principal component approach) could be implemented, but it would need to be tailored to each stimulus/task being considered. Any realistic attempt to leverage the convergence of receptor input performed by higher sensory areas would be a major undertaking that could not be simply added to this study. We also use a simple measure of response strength (peak-to-trough firing rate), and although it is likely one of the key elements of the response, other aspects could be considered. Particularly, synchrony among receptors has been shown to encode frequency modulations that occur during communication (Benda et al., 2006; Metzen et al., 2020). Changes in synchrony may occur as the stimulus strength changes and encode information that could enhance the detection and localization accuracy. Further experiments are required to better understand the importance of population synchrony in this context. Other neural mechanisms could be present and enhance the system’s sensitivity. We know, for example, that the presence of negative correlations in inter-spike intervals suppresses noise at low frequencies and could enhance the coding of low-frequency stimuli (lower than the frequency we use in this paper; (Chacron et al., 2001). For this reason, we propose that our sensitivity estimates are conservative, serving as a starting point for estimating the limits in detection and localization abilities.

Our analysis suggests that reliable detection or localization would require the integration of signals over a certain period by higher brain areas. This is a realistic perspective, as behavioral performance in various systems is more accurate for ongoing stimuli than for brief ones(Dizon and Litovsky, 2004; Gai et al., 2013). For example, estimates of the direction of motion in a “random dot display” integrate over time in the visual system of primates to reach a reliable decision after seconds of attending the stimulus (Ditterich et al., 2003; Kim and Shadlen, 1999). Spatial memory and navigation in sharks and turtles involve prolonged signal integration (Fagan et al., 2013). Furthermore, long-range search behavior in Diptera also relies on dynamic integration of spatial information (Kaushik et al., 2020), and directional coding in insect brains underscores the broad applicability of spatial integration strategies (Beetz & el Jundi, 2023). For weakly electric fish, it is not unrealistic to assume that the stimulus can be integrated over several hundred ms to support accurate detection and localization. However, it is not clear that this integration could occur over tens of seconds, particularly when relative movement requires the location estimate to be updated frequently. As a first step, our analysis considered spatially fixed signals; however, it will be imperative in future studies to take into account both the spatial and temporal dynamics of the conspecific signals. According to this perspective, localization accuracy, and thus behavioral decisions, depends on the spatial dynamic during the interaction. Moreover, this spatial dynamic can be leveraged as a means to collect spatial information. Various behavioral strategies can contribute to localizing a stimulus. For example, when localization is difficult, movements toward the target increase signal strength and help confirm the position of the second fish (Fagan et al., 2013; Kaushik et al., 2020). Lateralization, rather than precise azimuth localization, can be used while a fish moves towards a target: if a stimulus is perceived as coming from the left or right, corrective turns realign the target. The individual would therefore move toward the target in a zig-zag pattern; this mechanism has been proposed to contribute to behavior in various systems (Beetz and el Jundi, 2023; Gerhardt et al., 2023; Pollack et al., 1984).

Detection could rely on the convergence of the whole population of receptors, thereby efficiently averaging out noise. Furthermore, the high receptor density rostrally improves detection for frontal azimuth because the EI hotspot is centered on this high receptor density area, leading to a high convergence of strong responses. Localization, however, must rely on a comparison of the EI strength across the body. When comparing the responses to stimuli from different azimuths, our analysis focuses on the neural responses that differ between the stimuli, weighing heavily the contribution of these neurons. This procedure to optimize the extraction of spatial information is essential because the EI might differ only slightly between two stimuli, and thus, the responses of most neurons will be identical across the stimuli locations. For localization, it is thus the convergence of the responses of a subset of neurons that differ in activation between the locations being compared that can support the accurate discrimination of azimuth. Surprisingly, we found that the high density of receptors on the rostral portion of the fish does not lead to a better discrimination of the frontal azimuth. This result could reflect the fact that the receptors on the head will have a stronger difference in response at the edge of the hot spot elicited by the sender moving across these receptor locations. We suggest that this would occur as the leading or trailing edge of the hotspot sweeps across the high-density areas. This could explain the modest increases in spatial resolution due to the increased rostral receptor density that we observed around 60° and 120°. This is in contrast with the contribution of the high density of receptors during prey capture (i.e., small objects), where localization accuracy is enhanced for the head and snout regions. The structure and convergence of receptors thus play different roles in the detection and localization of objects and conspecifics: while high density and convergence contribute markedly to the spatial coding of objects, they mostly contribute to the detection accuracy of conspecific signals.

In other systems, regions of high receptor density are typically associated with high spatial resolution. It is the case of the foveal region of the retina and of high receptor density regions of the somatosensory system like the fingers or lips in humans (Catania and Catania, 2015; Dacey, 1994; Nakamura et al., 1998). In other systems faced with spatially diffuse signals, like the auditory or olfactory systems, the extraction of spatial information typically relies on the comparisons between a limited number of inputs (e.g., two ears) and high convergence of receptors is more tightly involved with the accurate detection, rather than localization, of the signals (Carr and MacLeod, 2010; Chapman, 1982; Okada and Toh, 2006; Schnupp and Carr, 2009). Our results on the electrosensory system suggest that this is a general principle guiding the relationship between signal structure and the organization of sensory arrays. We argue that for signals that are spatially diffuse, receptor density and convergence will benefit detection abilities, and spatial information is extracted by comparing inputs at different locations without relying on a higher number of inputs to enhance localization. For signals that are spatially localized, a topographic mapping system is advantageous, and localization accuracy directly depends on the spatial resolution of the input array. This perspective invites speculation about the evolutionary pressures that shape sensory receptor distributions. Spatially diffuse signals may drive sensory system evolution towards maximizing detection through receptor convergence, whereas localized signals might select for heightened spatial resolution. Since both types of signals are relevant in the electrosensory system of knifefish, it would be shaped by different evolutionary pressures that do not automatically require the same optimal solution. Investigating these evolutionary implications could further illuminate sensory processing adaptations across diverse ecological niches. The electrosensory system provides a powerful way to compare these systems and reveal organizing principles because it processes both localized and diffuse signals for which receptor organization and convergence play distinct roles.

## Acknowledgments

The authors would like to thank Dr. Alexandra Schmidt for assistance with fluorescence microscopy. Note that a previous version of this paper was included in the PhD thesis of Ramachandra (2023) and Milam (2023)

## SUPPLEMENTARY INFORMATION

**Table S1:** Model parameters used for the prototypic seed neuron (see Methods for details).

| Description | Name | Value | Description | Name | Value |
| --- | --- | --- | --- | --- | --- |
| Membrane time constant (s) | $\tau_m$ | $15 \cdot 10^{-4}$ | Noise strength (A) | $A_\sigma$ | $15 \cdot 10^{-9}$ |
| Lean reversal potential (V) | $E_m$ | $-70 \cdot 10^{-3}$ | Adaptation reversal potential (V) | $E_a$ | $-80 \cdot 10^{-3}$ |
| Membrane resistance ($\Omega$) | $R_m$ | $15 \cdot 10^5$ | Adaptation increment (V) | $\Delta_a$ | $14.5 \cdot 10^{-8}$ |
| Spiking threshold (V) | $V_T$ | $-49 \cdot 10^{-3}$ | Adaptation time constant (s) | $\tau_a$ | $50 \cdot 10^{-3}$ |
| Reset potential (V) | $V_R$ | $-70 \cdot 10^{-3}$ | EOD amplitude (A) | $A_{EOD}$ | $1.7 \cdot 10^{-7}$ |
| Refractory period (s) | $t_R$ | $9 \cdot 10^{-4}$ | EOD baseline bias (A) | $\beta$ | $3 \cdot 10^{-8}$ |

**Figure S1:**
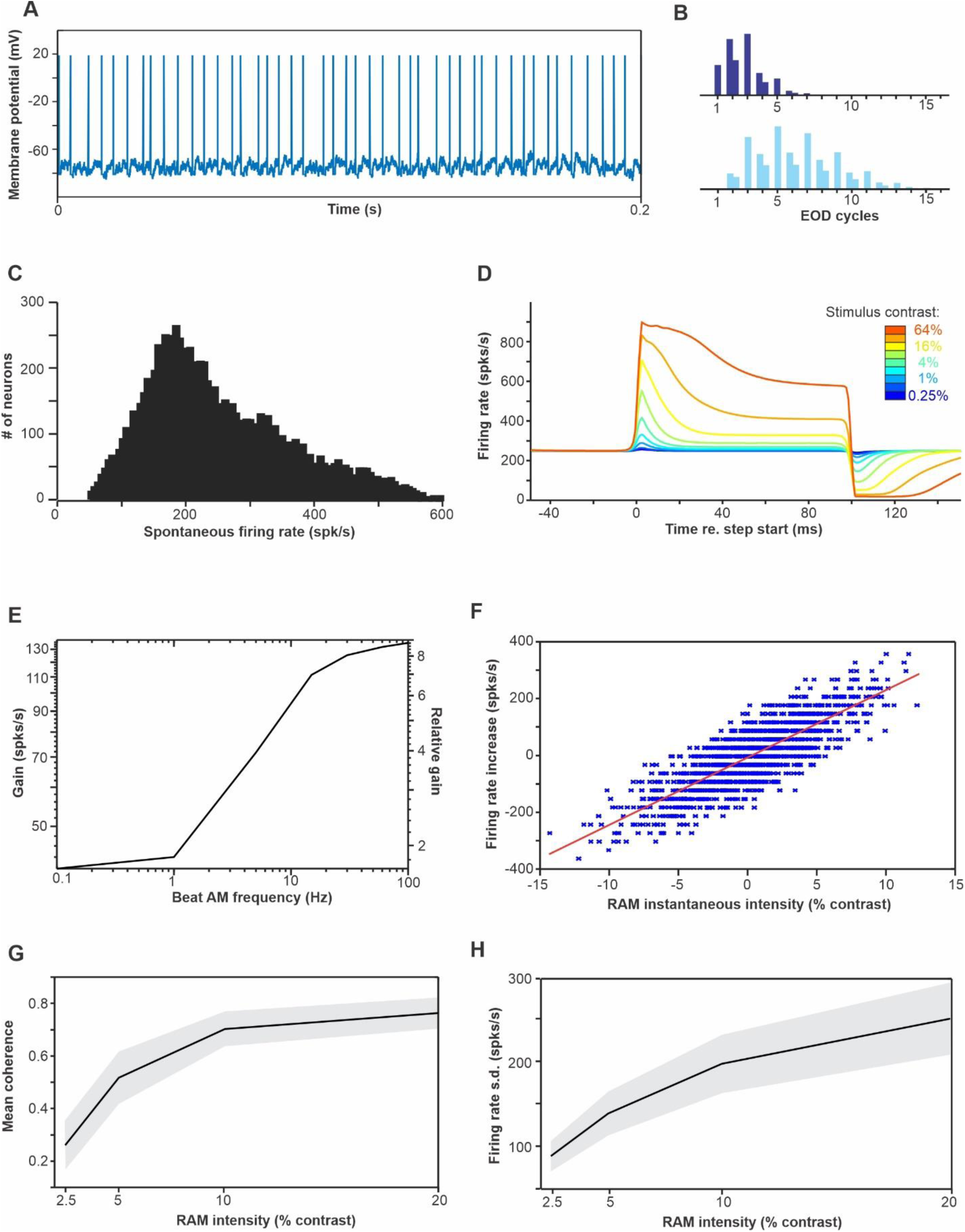
Average response properties of the heterogeneous population of modeled receptors. **A.** an example of the membrane potential and spiking pattern of a model response (spontaneous activity). **B.** Inter-spike intervale histogram of 2 different model neurons (spontaneous activity) showing phase locking to the EOD period and different firing tendencies. Note that the x-axis is expressed in multiples of the EOD period but since we used an EOD frequency of 1,000 Hz for simplicity, this also corresponds to ms. **C.** Distribution of spontaneous firing rate across our entire population. This distribution was achieved by selecting 26 seed neurons with firing rates unevenly distributed along this range and diversifying model parameters based on these seeds (see Methods). This range and distribution replicates published data (Bastian, 1981; Grewe et al., 2017; Ratnam and Nelson, 2000). In particular, the mean spontaneous firing rate was 251 spk/s with a CV of 0.45. **D.** Responses to step increases in EOD intensity. The strength of the peak response, the steady-state response, and the adaptation time course were matched to published data (Benda et al., 2005). **E.** Response gain to beat stimuli of different AM frequencies. Although we did not explore systematically the response of our model at different beat frequencies in the results section, we calculated the gain for a range of AM frequencies. This analysis helps us to evaluate the sensitivity of the neurons (see the absolute scale on the left) and it also helps to assert that the adaptation dynamic replicates some of the tuning properties of the neurons (see the relative scale on the right; gain for an AM of 1 Hz is normalized to 1). This average gain curve is comparable to experimental data (Chacron et al., 2005; Nelson et al., 1997). **F.** Relative firing rate during random amplitude modulations. We replicated a published analysis of receptor sensitivity (Gussin et al., 2007a) that measures the relative firing rate (relative to average) in successive 32 ms windows during the response to a low frequency (0 to 4 Hz) random amplitude modulation. We note that our scale is different from theirs: our ± 5% values correspond to the absolute contrast of the stimulus (i.e. s.d. of 10% contrast) whereas the scale in their Fig 3 has ± 50% being the min-to-max of their 10% contrast stimulus. When converted to the same scale, we find a gain slope of 23.7 spk/s/% (± 5.8 s.d.) comparable to the 17.7 spk/s/% (range 3.2 to 40.2) they found. **G.** Mean coherence at 10-30 Hz for random amplitude modulations (0-100 Hz) of different overall intensities (contrast). We qualitatively matched the coherence to values found in previous publications (Chacron et al., 2005; Grewe et al., 2017). **H.** Firing rate modulation in response to random amplitude modulations (0-300 Hz). We replicated the analysis in Grewe et al. (2017; see their figure S1A) which plots the standard deviation of the response (averaged over trials) as a function of stimulus contrast. Their population averages go from approximately 100 spk/s at 2.5% contrast to 200 spk/s at 20% contrast with a large variability among the population; we have a population average between 90 spk/s and 250 spk/s respectively.

**Figure S2:**
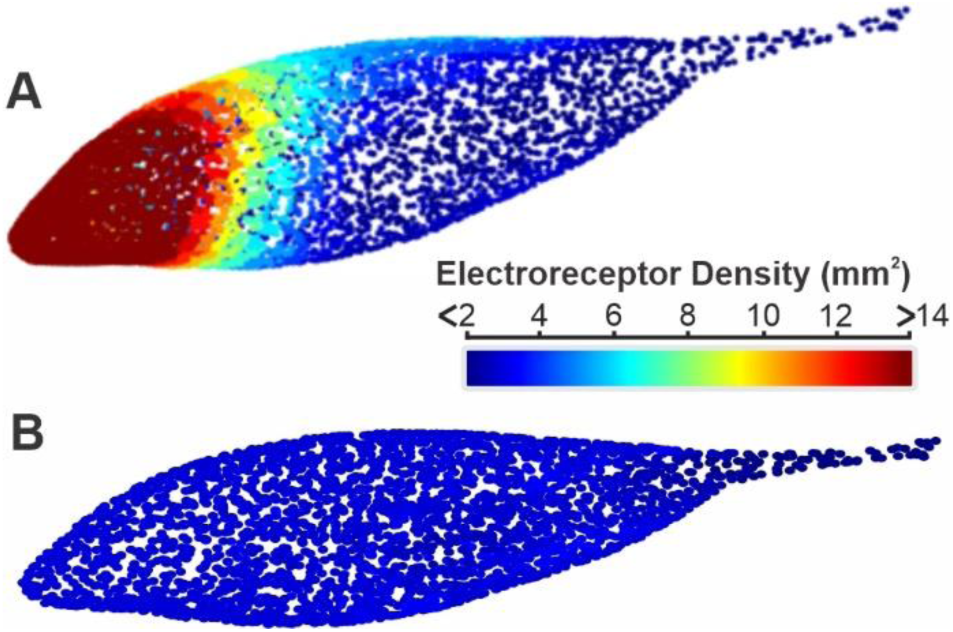
Receptor density compared between our full population (A) and our uniformly low-density population. **(B)**. Each dot shows the position of a receptor and the color reflects the density of receptors at this location. The uniform population was created by selecting, for each face of the mesh model, a subset of receptors from the full population to match the density of 2 receptors per mm^2^.

**Figure S3:**
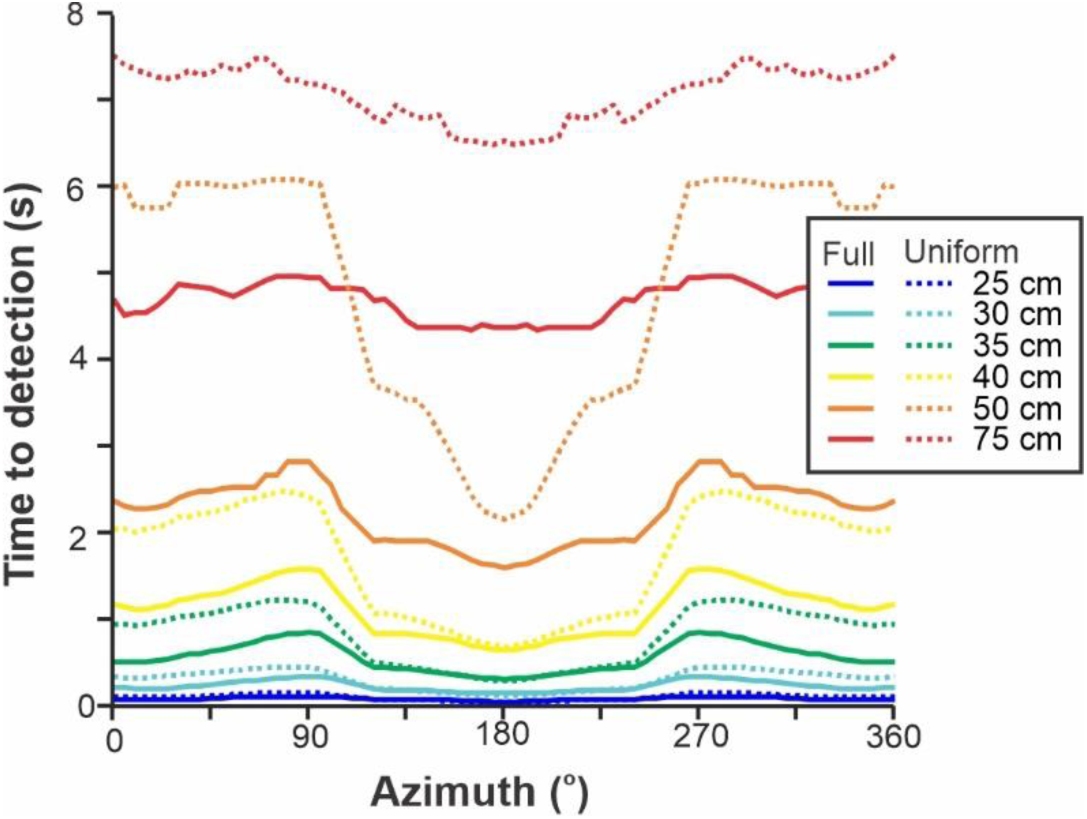
Detection performance as a function of source location and receptor structure. We compare a full population of receptors that includes a high density in rostral regions with a population that has a uniform density across the body matching the low density of the caudal region of the body (see Figure S2B). We present here the results for distances above 25 cm where we can see differences across azimuth and population structures. Our decoder considers that the response strength of receptors can be integrated across time and thus more noisy, weak responses require integration across longer periods to reach a reliable detection performance. We plot here the integration time required for our decoder to reach 95% detection accuracy and plot this here as a function of stimulus location.

